# Dissociating representations of object shape, real-world size, and mobility in human visual cortex

**DOI:** 10.64898/2026.07.05.736560

**Authors:** Simen Hagen, Yuanfang Zhao, Hans Op de Beeck, Marius V. Peelen

**Author notes:** Corresponding authors: Simen Hagen Marius Peelen Thomas van Aquinostraat 4, 6525 GD, Nijmegen, Netherlands. Author note: Simen Hagen; Yuanfang Zhao; Hans Op de Beeck, Marius V. Peelen. The authors declare no conflict of interest. Author contributions: SH: conceptualization and design, data collection, analyses, writing. YZ: conceptualization, writing. HOB: writing. MP: design, conceptualization, writing.

## Abstract

Object representations in the human ventral occipitotemporal cortex (VOTC) are organized along multiple dimensions, including shape (rectilinear vs. curvilinear), real-world size (large vs. small), and mobility (stationary vs. mobile). However, these dimensions are strongly correlated in naturalistic vision, making their separate contributions to VOTC organization unclear. For example, large objects (e.g., a wardrobe, a house) are typically rectilinear and stationary, while small objects (e.g., a ball, a cup) are more curvilinear and mobile. Here, we used fMRI, together with a new stimulus set that orthogonally manipulates shape, size, and mobility, to investigate the separate influences of these dimensions on VOTC organization. Example stimuli include air balloon (large, curvilinear, mobile), radar dish (large, curvilinear, stationary), and mailbox (small, rectilinear, stationary). Contrasts revealed that large (vs. small), rectilinear (vs. curvilinear), and stationary (vs. mobile) dimensions all independently evoked strong and overlapping activity in medio-anterior VOTC. This overlapping activity was at the intersection of the parahippocampal place area (PPA) and the ventral place-memory area (VPMA). Similar results were found at the intersection of the scene-selective occipital place area and the lateral place-memory area (LPMA). Finally, large (vs. small), but not rectilinear (vs. curvilinear) or stationary (vs. mobile) activity, was found in additional posterior ventral scene-selective regions, as well as in early visual cortex. Overall, these results indicate that object shape, real-world size, and mobility dimensions all independently activate scene-selective PPA and OPA, showing joint selectivity for distinct low- and high-level object properties that are highly correlated in naturalistic vision.

## Introduction

The human ventral occipitotemporal cortex (VOTC) is critically involved in visual object recognition (Farah, 2004) and contains a systematic and consistent functional organization for object categories (Grill-Spector & Weiner, 2014). However, the factors that determine the organization remain debated.

One major organizing principle is real-world object size, associated with a medio-lateral dissociation in the VOTC, where large (vs. small) and small (vs. large) objects evoke strong medial and lateral activity, respectively (e.g., Konkle & Oliva, 2012a; Konkle & Caramazza, 2013). This real-world size organization may reflect the different functional properties of large and small objects, such as their graspability (Bainbridge & Oliva, 2015) or their usefulness as landmarks for navigation (Epstein & Vass, 2014). However, real-world size correlates with visual properties that could alternatively or additionally account for this organization. For example, large and small objects covary with rectilinearity, the former being more rectilinear and the latter more curvilinear (e.g., Long et al., 2016), and simple rectilinear and curvilinear shapes evoke strong responses in medial and lateral VOTC, respectively (Nasr et al., 2014; Yue et al., 2020). Indeed, unrecognizable large and small objects that maintain their differences in low-level visual properties generate a similar large scale medio-lateral organization as large versus small recognizable objects (Long et al., 2018). This has led to the proposition that the real-world size organization could be driven largely by low-level visual properties that covary with a higher-level conceptual difference (e.g., Long et al., 2018).

This issue has been examined in more detail in the scene-selective parahippocampal place area (PPA; Epstein & Kanwisher, 1998; Aguirre et al., 1998), a region in medial VOTC that responds strongly to large (vs. small) manmade objects (e.g., Konkle & Caramazza, 2013). The PPA responds to several visual properties covarying with large object size (Hasson et al., 2003; Nasr et al., 2014; Rajimehr et al., 2011), in line with a visual feature account. Interestingly, however, the PPA also responds to large (vs. small) objects in congenitally blind participants (He et al., 2013), suggesting that large-object selectivity cannot be fully explained by visual feature differences. Instead, the PPA may reflect the degree to which objects evoke a sense of space (Mullally & Maguire, 2011) or the degree to which they are useful for navigation (Aguirre et al., 1998; Epstein & Vass, 2014). Large objects, particularly those that are stationary (e.g., buildings, large furniture), are most effective in defining local space (Mullally & Maguire, 2011) and are also most useful as landmarks for navigation. Accordingly, the PPA responds most strongly to objects that are large and stationary (Mullally & Maguire, 2011; Troiani et al., 2014). Importantly, however, previous work has not experimentally manipulated size and mobility as separate factors, leaving open the possibility that the key object property driving PPA is being stationary rather than being large. Another possibility is that the PPA responds selectively only to objects that are both large and stationary (e.g., buildings), rather than to either of these properties separately. Finally, it is possible that the effects of size and mobility are largely accounted for by covarying shape features, with large and stationary objects having more rectilinear features.

Here we used fMRI, together with a new stimulus set that orthogonally manipulates shape (rectilinear, curvilinear), size (large, small), and mobility (stationary, mobile) (Fig. 1), to investigate which of these factors are reflected in VOTC. The critical aspect of this design is that it allows us to examine rectilinear (vs. curvilinear), large (vs. small), and stationary (vs. mobile) object responses independent of each other. We do this within functionally-localized PPA, as well as other scene-selective regions. Moreover, we examine the spatial distribution of each of the properties separately across the VOTC, thereby testing the large-scale organization of size, shape, and mobility.

**Figure 1.**
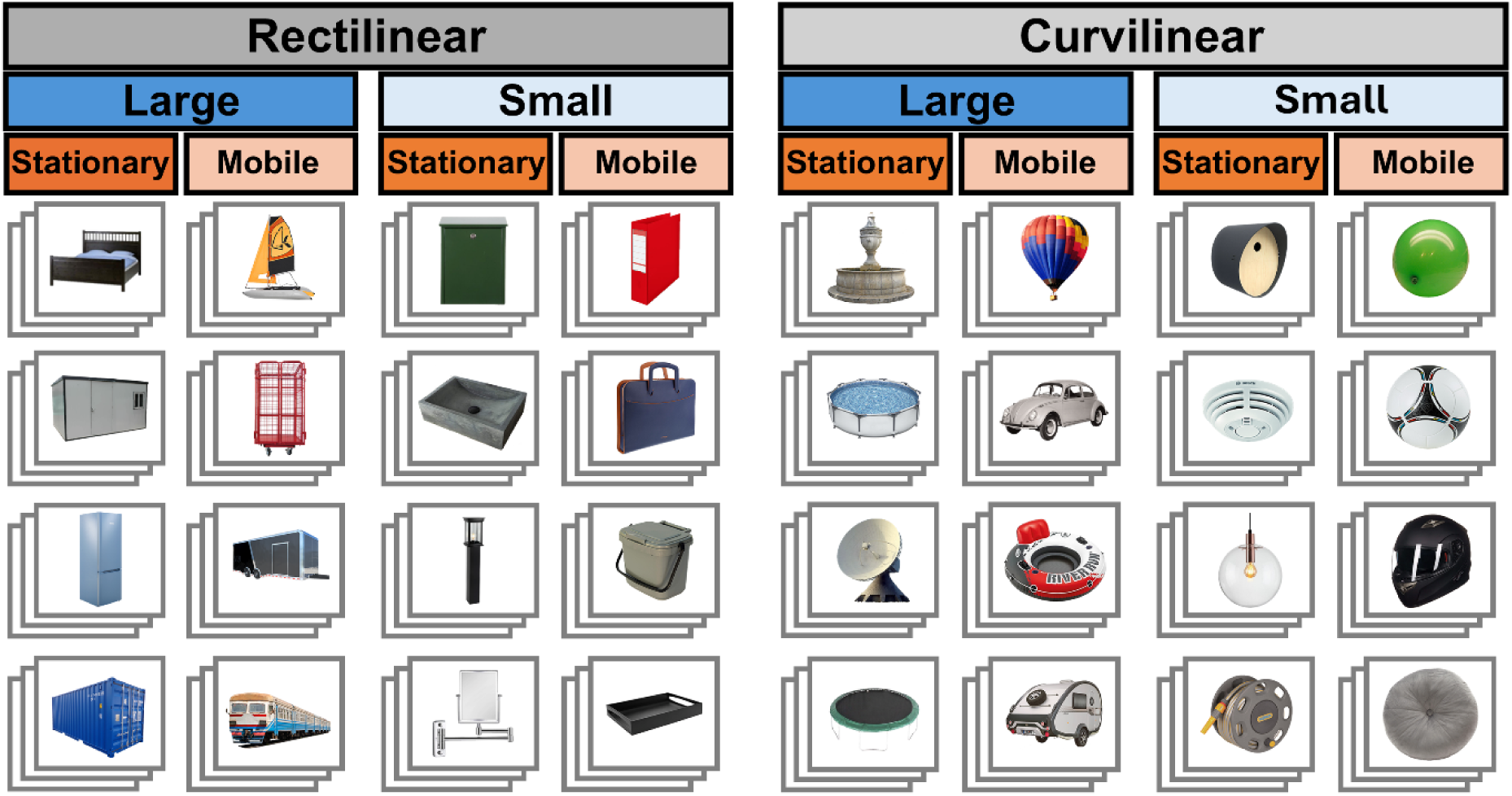
Experimental design and stimuli. A two (size: large, small) by two (shape: rectilinear, curvilinear) by two (mobility: stationary, mobile) design was used to examine independent size, shape, and mobility fMRI responses. The eight experimental conditions were composed of four object categories each with three different image exemplars.

## Methods

### Participants

Twenty-two participants (*M* = 22.3 years; *SD* = 3.2 years; age range: 18 - 30; females = 11, males = 11) volunteered to participate in return for course credit or monetary compensation. The participant sample was set to match that of Konkle & Oliva (2012a, *N* = 22). The participants were recruited through the institutes’ internal recruitment system. Informed consent was obtained prior to the experiment. The experiment was approved by the local ethics committee (CMO Arnhem-Nijmegen). All data were collected in the years 2022 and 2023.

### Stimuli

The experimental stimuli were selected for a two (object shape: rectilinear, curvilinear) by two (real-world object size: large, small) by two (real-world object mobility: stationary, mobile) design. This design yielded eight conditions: rectilinear-large-stationary, rectilinear-small-stationary; rectilinear-large-mobile, rectilinear-small-mobile; curvilinear-large-stationary, curvilinear-small-stationary; curvilinear-large-mobile, curvilinear-small-mobile (**Fig. 1**). Each condition was represented by four object categories with three different object exemplars each (**Fig. 1**; full list of object categories in **Table S1**). The object images were found online, cropped from their background, and pasted on a phase-scrambled background. The phase-scrambled background was different for each object exemplar and created in the following way: (1) randomly select one image exemplar from an object category from the other conditions, (2) phase scramble this image, (3) paste the object exemplar on top of the phase-scrambled object. This procedure was done to control the conditions on low-level properties of color, spatial frequency, and luminance (see stats for these low-level properties below). The background subtended 20.9° [H] x 15.7° [V] visual angle (1200 x 900 pixels), while the centered objects subtended 4.35° visual angle (250 pixels) in the longest dimension of the object.

The main contrasts (large vs. small; rectilinear vs. curvilinear; stationary vs. mobile) did not differ in terms of spatial frequencies (*p*s > .13), color in both the RGB and LAB space (*p*s > .05), Luminance (L dimension in LAB space) (*p*s > .3; see Bainbridge & Oliva (2015) for a description of the procedures for statistically testing these properties). As expected, the shape contrast, but not the size and mobility contrasts, differed on rectilinearity (*p* < .001, *p* > .980, *p >* .271, respectively; see Li & Bonner (2020) for a description of the toolbox used to compute rectilinearity). The stimuli for the conditions reflected in the two-way interaction also did not differ in terms of spatial frequencies (*p*s > .05), color in both the RGB and LAB space (*p*s > 0.129), and luminance (L dimension in LAB space) (*p*s > .284). Moreover, the shape conditions (e.g., large & rectilinear vs. large & curvilinear), but not the size and mobility conditions (e.g., rectilinear & large vs. rectilinear & small), differed in rectilinearity (*ps* < .001, *p*s > .574, *p*s > .092, respectively).

The localizer stimuli consisted of objects from scenes (N=20), large objects (N=60), small objects (N=60), buildings (N=8) and boxes (N=8). The buildings and boxes were not further analyzed. The small and large objects subtended 4.35° visual angles (250 pixels) in the longest dimension of the object, while the scenes subtended 4.35° visual angles in both horizontal and vertical dimensions.

### Experimental design

In the experimental task, the participants viewed object stimuli one-by-one for 500 milliseconds (msec) each, presented in a mini block design, with 32 mini-blocks per run, each showing six images from the same object category (e.g., air balloon). Each image and mini block were interspersed by a fixation cross for one and two seconds, respectively, and each run was flanked by a fixation cross for 10 seconds. There was a total of four runs each showing the same object stimuli with the order of mini-blocks randomized. Participants performed an orthogonal task by pushing a response button every time there was a repetition of an object (e.g., the same air balloon presented twice in a row). There were four randomly determined repetitions per condition (e.g., rectilinear-large-stationary).

For the localizer task, the design was equivalent to the experimental task design, with the exception that each run consisted of 30 mini blocks, each showing 11 images from the same object category, and an inter-stimulus-fixation-cross interval of 300 msec. The participant also completed four additional experimental runs that were part of a different experiment and that are not analyzed and reported here.

### fMRI acquisition, preprocessing, and analyses

#### Acquisition

The fMRI data was acquired with a Siemens 3T MAGNETOM Skyra scanner (Siemens AG, Healthcare Sector, Erlangen, Germany) using a 32-channel head coil. At the start of each experimental session, a high-resolution T1-weighted anatomical image was acquired with a MPRAGE sequence (TR 2300 msec, TE 3 msec, flip angle: 8°, 1 x 1 x 1 mm isotropic voxel size, 192 sagittal slices (X), 256 coronal slices (Y); FoV 192 mm x 256 mm (Y)). Subsequently, whole-brain functional blood oxygenation level-dependent (BOLD) activity was acquired using a T2*-weighted gradient echo EPI sequence with 4x multiband acceleration factor (TR 1500 msec, TE 33 msec, flip angle 75°, 2.048 mm x 2.048 mm x 2.00 mm voxel size, FoV: 212.99 mm x 212.99 mm, 68 slices. Each experimental and localizer run acquired 232 and 272 volumes, respectively.

#### Preprocessing

The Functional Magnetic Resonance Imaging of the Brain Software Library (FSL, Oxford, UK) was used for data preprocessing. The first 3 volumes of each run were removed. The subsequent preprocessing steps included a field-map correction, a two-step spatial realignment of the functional images, high pass filtering (100 Hz cutoff), spatial smoothing (5 mm FWHM gaussian kernel), normalization to MNI 152 space, and independent component analysis (ICA) to remove head motion.

#### Analysis

Using FSL (FSL, Oxford, UK), separately for experimental and localizer runs, each condition was modeled as a convolution of a boxcar with a human hemodynamic response function. Contrasts were computed at the run-level and the run-based values were averaged separately for each participant and condition. Group-level statistical maps were created from effect and variance maps. Custom scripts in Python were used for region of interest (ROI) analyses, which were conducted in volumetric space. ROIs were created by intersecting independent localizer data (see Results section; **Fig. 2A**) with probabilistic masks. Intersection with probabilistic masks were used to localize the PPA, occipital place area (OPA), and retrosplenial complex (RSC) (Zhen et al., 2015). The probabilistic thresholds of the masks (and thus the size) were set to capture at least 50 scene-selective voxels per subject (PPA: *p* > 0.4; OPA: *p* > 0.1; RSC: *p* > 0.5). All probabilistic masks are visualized on the cortical surface in Figures 2 and 4. Group-level visualizations of brain maps were done in fsaverage surface space using Nilearn (Python).

**Figure 2.**
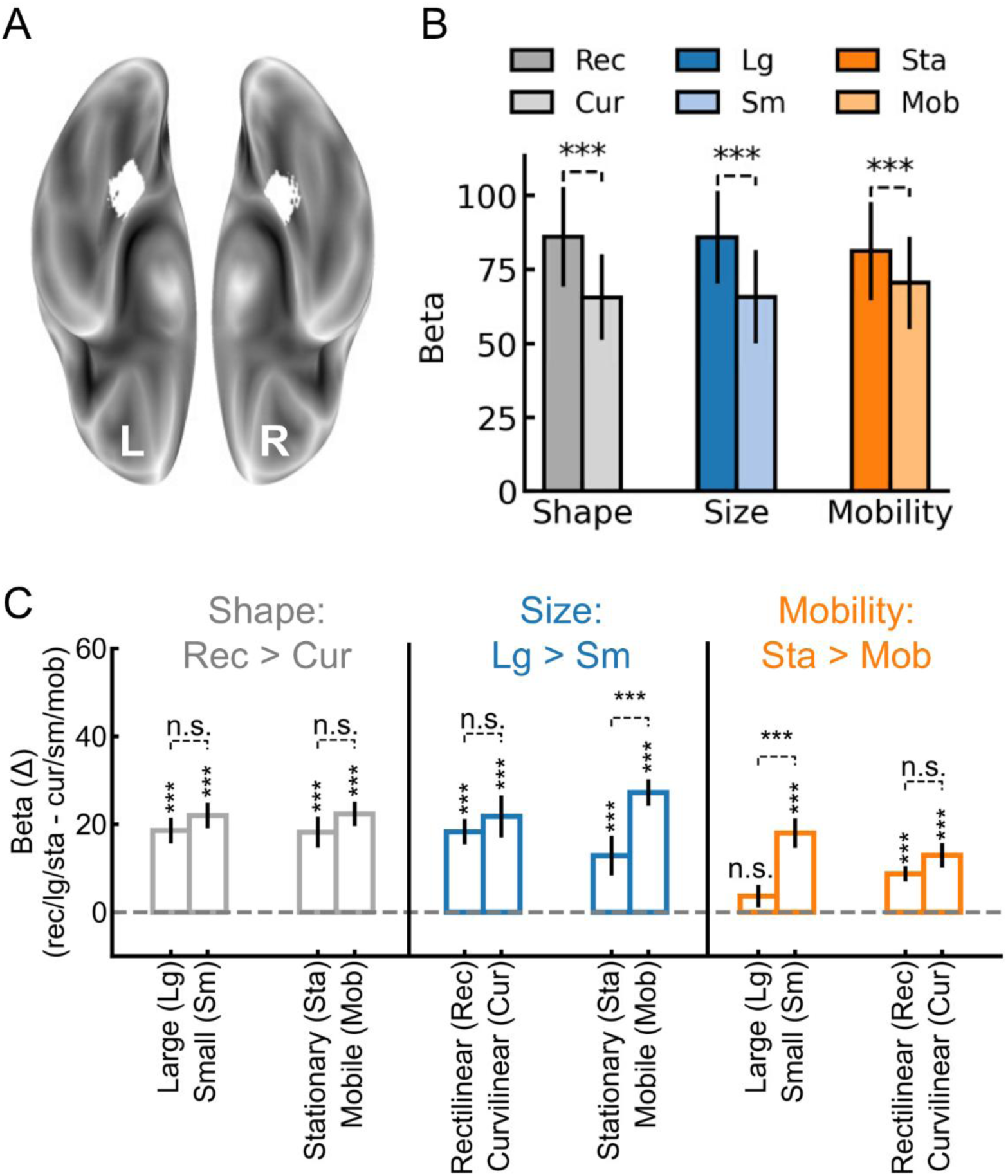
Responsiveness of the PPA to objects varying in shape (rectilinear, curvilinear), real-world size (large, small), and mobility (stationary, mobile). (**A**) In a region-of-interest (ROI) analysis, the PPA was defined by selecting the 50 most scene-selective (scene > small objects) voxels within a bilateral probabilistic PPA mask. (**B**) The beta values to shape (rectilinear, curvilinear), size (large, small), and mobility (stationary, mobile) conditions were extracted separately for each participant. Note that for each property (e.g., size), the other two properties (e.g., shape and mobility) are balanced. (**C**) The selective response was computed for each property (e.g., lg-sm) separately for each other object property (e.g., rectilinear, curvilinear, stationary, mobile). A positive value indicates a selective response in the direction of the preferred property (i.e., lg > sm; rec > cur; sta > mob), while a negative value would indicate a selective response in the direction of the non-preferred property (i.e., lg < sm; rec < cur; sta < mob). Error bars represent 95% confidence intervals (CI). N.s. and *** represent p > 0.05 and p < 0.001, respectively.

##### Missing data

One participant had three instead of four experimental runs due to a technical error with the event timing file.

## Results

### 1. Rectilinear, large, and stationary object properties independently activate the PPA

We first examined whether the PPA responds independently to shape (rectilinear, curvilinear), real-world size (large, small) and mobility (stationary, mobile) properties. In a targeted region-of-interest (ROI) analysis, we independently localized the PPA by selecting for each subject the 50 most scene-selective voxels with the restriction that they had a Z-score > 1.645 (*p* < 0.05, one-tailed) (vs. object; independent localizer runs) within a bilateral probabilistic PPA mask (mask visualized on the cortical surface in **Fig. 2**). All participants had at least 50 voxels with Z > 1.645 scene-selective activity. The beta values for each of the eight experimental conditions (e.g., rectilinear-large-stationary) were extracted within the PPA masks and submitted to a repeated measures analysis of variance (ANOVA) with shape (rectilinear, curvilinear), size (large, small), and mobility (stationary, mobile) as within-subjects factors. Given our study design where shape, real-world size, and mobility are orthogonal, the main effects of the ANOVA examine whether the PPA responds to each property when controlling the other properties. The significant main effects showed that the PPA responded independently to shape (rectilinear > curvilinear), *F*(1,21) = 54.31, *p* < 0.001, size (large > small), *F*(1,21) = 46.57, *p* < 0.001, and mobility (stationary > mobile), *F*(1,21) = 39.66, *p* < 0.001 (**Fig. 2A**). Thus, the PPA is activated by objects that are rectilinear, large, or stationary.

We further probed the degree of independence between the three properties in the PPA by testing for interactions between them. There was a significant three-way interaction between shape, size, and mobility, *F*(1,21) = 13.81, *p* = 0.001. We followed this up by testing for two-way interactions. There were no interactions between shape and size, *F*(1,21) = 0.43, *p* = 0.522, and shape and mobility, *F*(1,21) = 1.74, *p* = 0.202 (**Fig. 2C**), showing that rectilinear (vs. curvilinear) did not differ across size (large, small) and mobility (stationary, mobile), and that large (vs. small) and stationary (vs. mobile) did not differ across shape (rectilinear, curvilinear). In contrast, size interacted with mobility, *F*(1,21) = 8.59, *p* = 0.008, reflecting that large (vs. small) was lower for stationary (but significant relative to 0: *t(21)* = 2.87, *p* = 0.009) than mobile objects, and that stationary (vs. mobile) was smaller for large (and non-significant relative to 0: *t*(21) = 1.41, *p* = 0.174) than small objects. Thus, while the effects of size and mobility were each independent of shape, the effects of size and mobility interacted.

Based on the significant two-way interaction between size and mobility, and previous findings that the combination of large and stationary drives PPA activity (Mullally & Maguire, 2011), we examined whether large-and-stationary objects differed from large-and-mobile, small-and-stationary, and small-and-mobile objects. Large-and-stationary objects did not differ from large-and-mobile objects, *t*(21) = 1.41, *p* = 0.174, while both large conditions evoked stronger activity than both small conditions, *t*s(21) > 2.85, *p*s < 0.01, and small-and-stationary objects evoked stronger activity than small and mobile objects, *t*s(21) < 5.37, *p* < 0.001 (**Fig. 3B**). Thus, the PPA showed a preference for large and stationary properties, but not specifically their combination.

**Figure 3.**
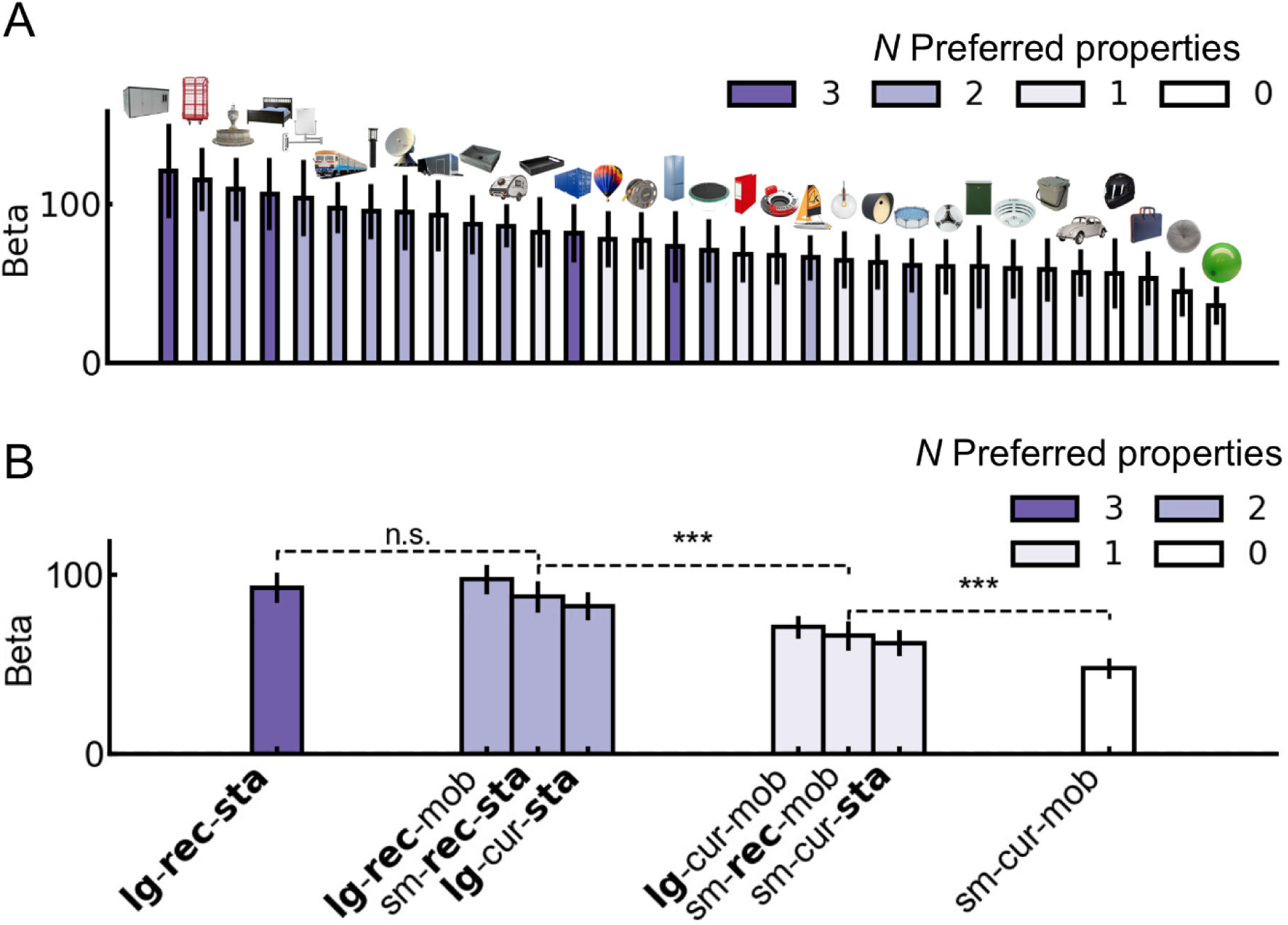
Responsiveness of the PPA to objects varying in the number of PPA-preferred properties. (**A**): Beta values for each object category, ranked from highest (left) to lowest (right) on evoked response. (**B**): Beta values for each of the eight experimental conditions, ranked from highest (left) to lowest (right) on number of preferred properties. Error bars represent 95% CIs. N.s. and *** represent p > 0.05 and p < 0.001, respectively. Bolded x-tick text indicates PPA-preferred properties. Abbreviations: lg: large, sm: small, rec: rectilinear; cur: curvilinear, sta: stationary, mob: mobile.

Finally, we examined whether PPA activity was influenced by the number of hypothesized PPA-preferred properties (i.e., large, rectilinear, stationary). At the object level, PPA activity generally increased as a function of the number of preferred properties (**Fig. 3A**). Thus, we aggregated the eight conditions as a function of number of preferred properties (e.g., sm-cur-mob objects have zero preferred properties, while lg-rec-sta objects have three preferred properties) to examine for a statistical difference between each step. While there was a significant increase from zero up to two preferred properties, *t*s(21) > 5.20, *p*s < 0.001, the response to objects with three preferred properties did not differ from the response to objects with two preferred properties, *t*(21) = 1.75, *p* = 0.095 (**Fig. 3B**). This is consistent with the above analysis of less independence among a subset of the properties (i.e., redundance across properties in driving the response).

### 2. Rectilinear, large, and stationary objects independently activate a broader scene perception network

We examined the independence of rectilinear (vs. curvilinear), large (vs. small), and stationary (vs. mobile) selective activity in other regions of a broader scene-selective network. First, we examined two cortical scene-selective regions, the occipital place area/transverse occipital sulcus (OPA/TOS) (Grill-Spector, 2003), and the retrosplenial complex (RSC) (Maguire, 2001). While these regions have been shown to respond to the properties investigated here (Auger & Maguire, 2012; He et al., 2013; Nasr et al., 2014), as for the PPA, they have not been investigated independently. Following the same procedure as used for the PPA, we intersected independent scene-localizer (scene > small manmade objects) data with bilateral probabilistic ROIs and selected the 50 most scene-selective voxels with the restriction of a Z score above 1.645 (masks visualized on the cortical surface in **Fig. 4**). All participants had at least 50 scene-selective voxels with Z > 1.645 (*p* < 0.05, one tailed) in the OPA and RSC. For the OPA, the significant main effects showed that it responded to shape (rectilinear > curvilinear), *F*(1,21) = 30.92, *p* < 0.001, size (large > small), *F*(1,21) = 58.53, *p* < 0.001, and mobility (stationary > mobile), *F*(1,21) = 26.87, *p* < 0.001 (Fig. 4A, right). For the RSC, the significant main effects showed that it responded to shape (rectilinear > curvilinear), *F*(1,21) = 17.22, *p* < 0.001, size (large > small), *F*(1,21) = 19.30, *p* < 0.001, but not mobility (stationary > mobile), *F*(1,21) = 1.74, *p* = 0.20 (Fig. 4A, right). The regions showed qualitatively similar responses as the PPA when considering the different combinations of preferred properties (**Fig. S1**). Thus, like the PPA, the OPA responded to all three properties (rectilinear, large, stationary), while the RSC responded to rectilinear and large objects but did not show a preference for stationary (vs. mobile) objects. Overall, this shows that multiple nodes of the scene-selective network selectively respond to rectilinear, large, or stationary objects.

**Figure 4.**
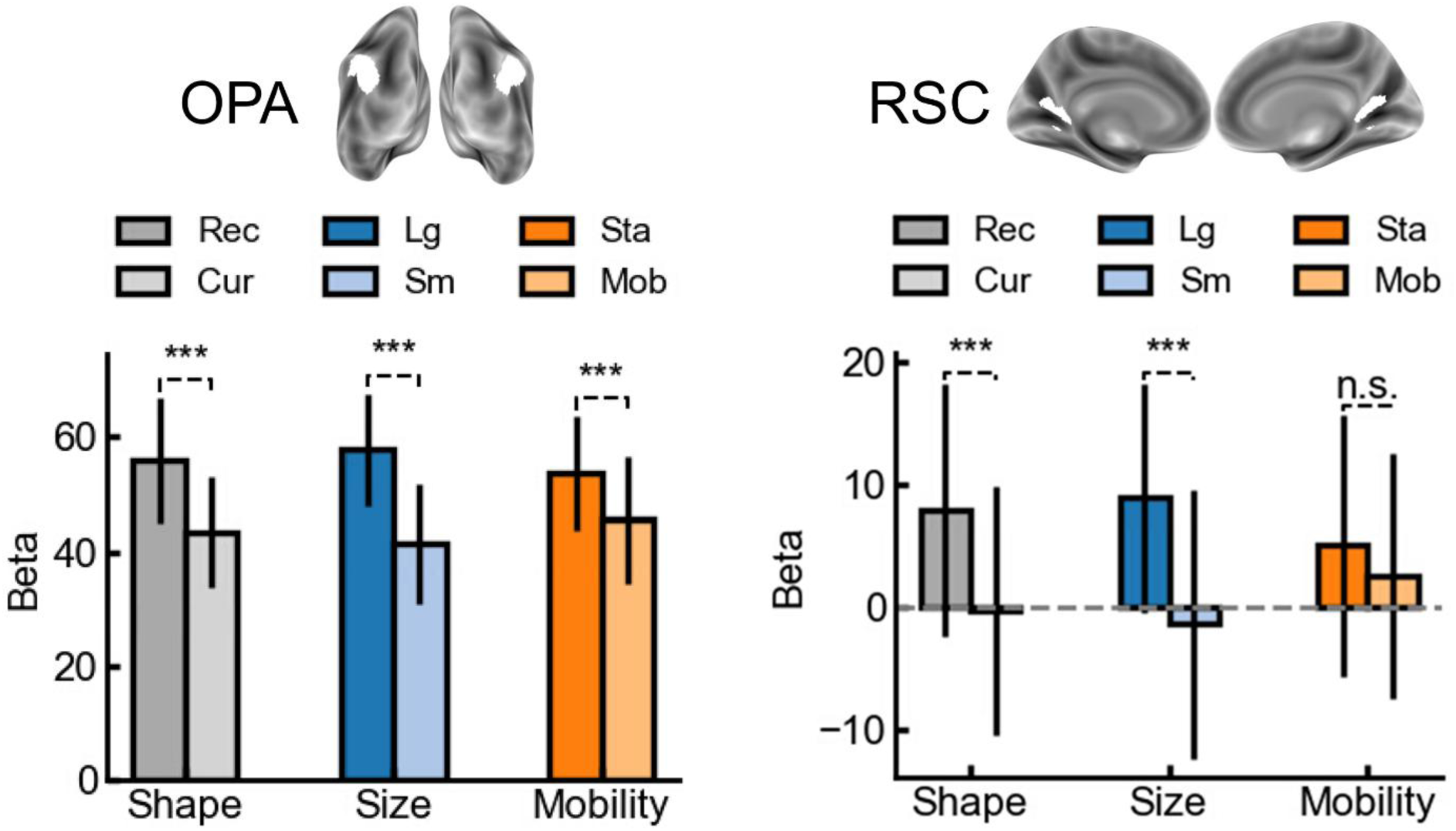
Responsiveness of the occipital place Area (OPA) and retrosplenial complex (RSC), to objects varying in shape (rectilinear, curvilinear), real-world size (large, small), and mobility (stationary, mobile). The beta values to shape (rectilinear, curvilinear), size (large, small), and mobility (stationary, mobile) conditions in OPA and RSC. Error bars represent 95% CIs. N.s., and *** represent p > 0.05, and p < 0.001, respectively.

### 3. Spatial distribution of overlapping rectilinear, large, and stationary activity in the VOTC

Our results indicate that rectilinear, large, and stationary object properties all drive PPA activity. Thus, we examined if this clustering is specific to PPA and other scene-selective regions rather than non-scene-selective regions. Specifically, we isolated all voxels in which rectilinear (vs. curvilinear), large (vs. small), and stationary (vs. mobile) showed activity, and plotted these “overlap” voxels on the cortical surface. Moreover, we superimposed probabilistic maps of scene-perception and -memory regions (Steel et al., 2021; 2025). We identified clusters of the “overlap” voxels (vertices on the cortical surface) on both the ventral and lateral surfaces (**Fig. 5**). Notably, on the ventral surface, the overlap voxels clustered in and around the PPA and ventral place-memory area (VPMA) (Fig. 5A, left), while on the lateral surface they clustered in and around the OPA and the lateral place-memory area (LPMA) (Fig. 5A, right). No other regions showed activity, even at the liberal threshold *p* < 0.05 (of Z > 2.3). Thus, the clustering of activity to all the three properties was remarkably specific to scene-selective regions.

**Figure 5.**
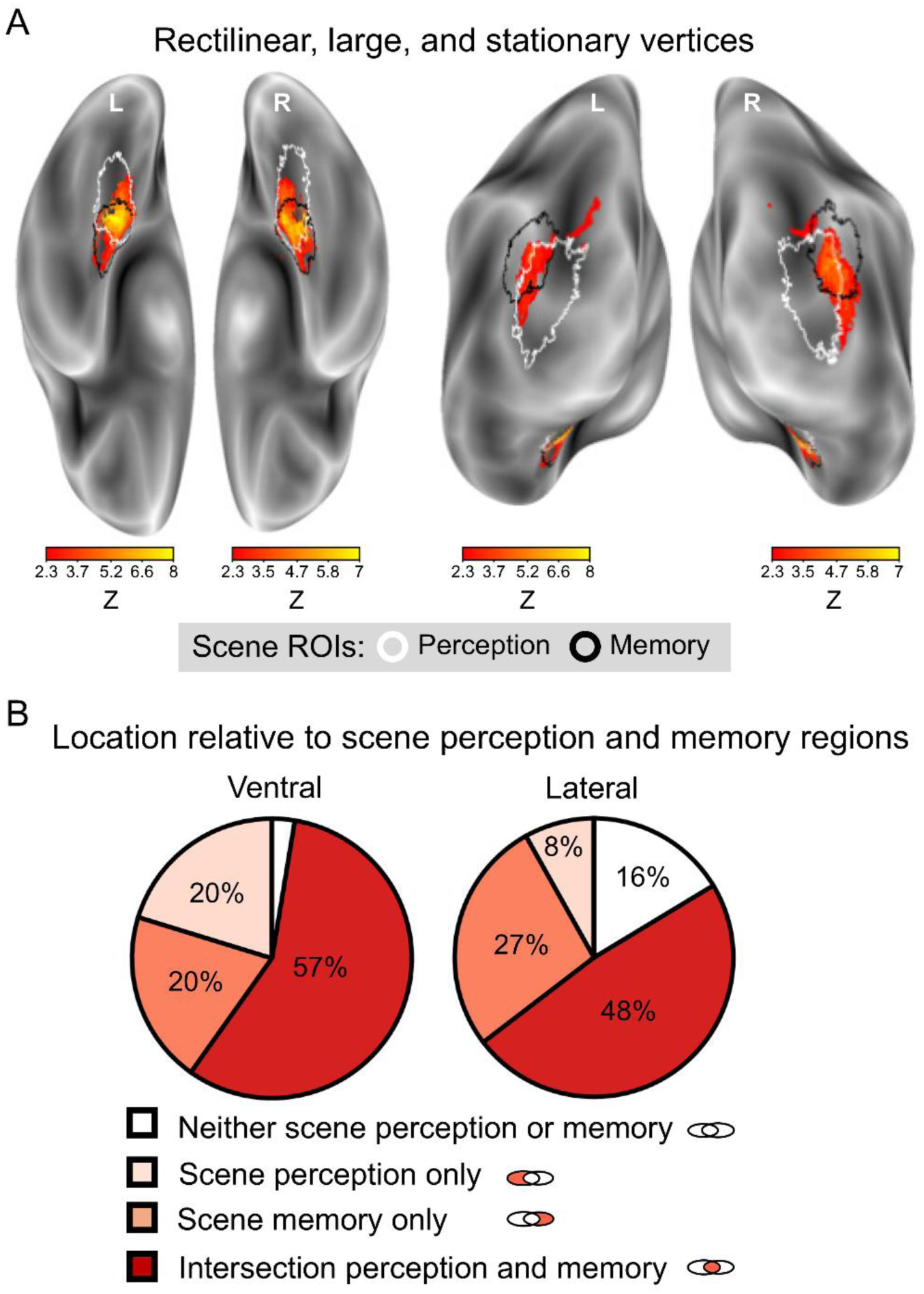
“**Overlap” voxels with rectilinear, large, and stationary selective activity plotted on the cortical surface.** (**A**) Voxels that are selective to all three properties at a threshold of Z > 2.3 (p < 0.05) projected on cortical surface maps. The color bar indicates the Z-score in the overlap voxels for the property with the lowest Z-score. White and black contours outline probabilistic scene-perception and - memory regions, respectively (Steel et al., 2025). (**B**) Percentage of “overlap” (selective to rectilinear, large, stationary objects) vertices (out of total “overlap” vertices).

We computed the proportion of overlap voxels (out of total overlap voxels) located within the perceptual and memory regions at Z > 3.1 (*p* < 0.001) (Fig. 5B). On the ventral surface, 97.38% was located within either the perceptual or memory regions (i.e., the regions collapsed), of which 57.20% was in the intersection between PPA and VPMA, 20.25% in unique PPA (most posterior), and 19.93% in unique VPMA (most anterior) (Fig. 5B, left). On the lateral surface, 83.65% of the overlap voxels were in either the perceptual or memory regions, of which 48.25% was in the intersection between OPA and LPMA, 8.12% in unique OPA, and 27.28% in unique LPMA (Fig. 5B right). Thus, the highest proportion of overlap voxels were in mid-anterior PPA, where scene perception and memory processes intersect.

### 4. Spatial distribution of separate rectilinear, large, and stationary activity across the VOTC

Finally, in exploratory analyses, we examined, across the whole brain, whether other regions were sensitive to the manipulated object properties. For example, shape and real-world size have both been associated with medio-lateral dissociations in the VOTC, with rectilinear (vs. curvilinear) and large (vs. small) evoking strong activity in the medial VOTC, while curvilinear (vs. rectilinear) and small (vs. large) evoking strong activity in lateral VOTC (Konkle & Oliva, 2012a; Nasr et al., 2014; Yue et al., 2020). Here, we could examine whether similar maps are found after controlling for covarying properties. Maps are plotted at a liberal threshold of *p* < 0.01 (one-tailed; Z > 2.3; cluster threshold *p* < 0.05) for visualization purposes.

All three properties showed consistent activation across participants in several regions of the VOTC (Fig. 6). For shape, we found rectilinear (vs. curvilinear) activity in expected medial ventral regions of the VOTC, with peak activity in the PPA, and on medial-dorsal regions of the lateral VOTC. Curvilinear (vs. rectilinear) activity was found in expected ventral lateral VOTC regions and lateral posterior VOTC regions. Thus, the shape maps showed the expected medio-lateral organization. For size, we found large (vs. small) activity in expected medial ventral regions of the VOTC, extending from the PPA to posterior ventral medial regions, and in medial dorsal regions. This is consistent with previous studies, and with an independent localizer of large and small objects (not controlling shape and mobility) in our own data (**Fig. S2**). However, we did not find the expected small (vs. large) activity in ventral lateral regions, or in the posterior lateral regions. For mobility, we found expected stationary (vs. mobile) activity on the ventral medial surface in vicinity of the PPA, and in dorsal posterior regions in the vicinity of the OPA. However, we also found strong unexpected activity for stationary (vs. mobile) activity on the lateral surface in the vicinity of the lateral occipital complex (LOC). Moreover, we found mobile (vs. stationary) activity in the ventral early visual cortex, consistent with previous work (Mullally & Maguire, 2011). Thus, each property on their own also activated an extensive network that diverged from the scene-selective network, presumably reflecting their contribution to other behaviorally relevant functions.

**Figure 6.**
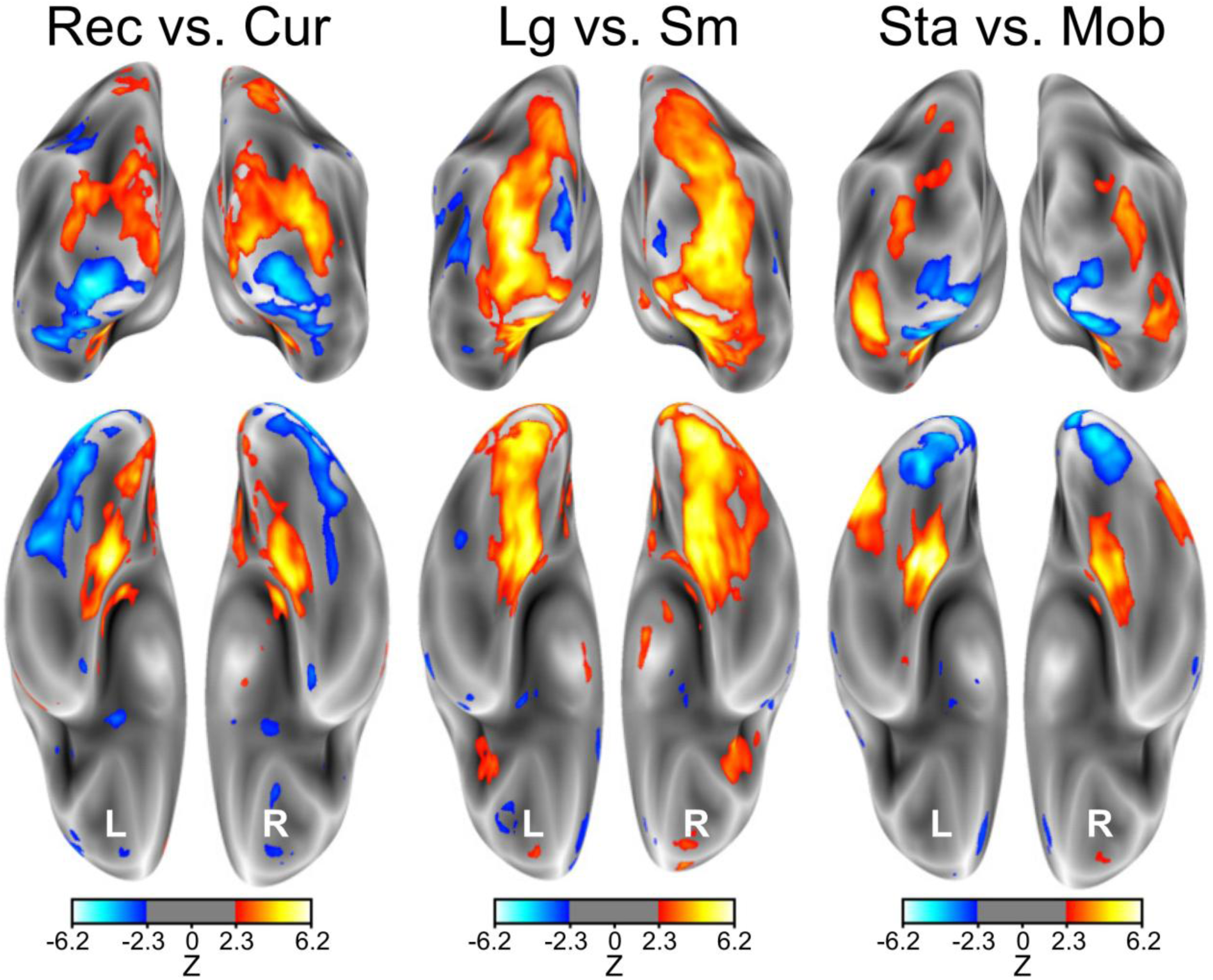
Whole-brain activity for shape (rectilinear vs. curvilinear), size (large vs. small), and mobility (stationary vs. mobile). Group-level (N=22) contrasts for shape (left), size (middle), and mobility (right). Positive values indicate stronger BOLD signal to rectilinear, large, and stationary objects. A liberal threshold of p < 0.01 (one-tailed; Z > 2.3) is used for visualization.

## Discussion

Real-world size is a salient object property that is automatically perceived from object images (e.g., Konkle & Oliva, 2012b; Hagen et al., 2024), is decoded from brain activity as early as 150 msec after stimulus onset (Khaligh-Razavi et al., 2018) and evokes a medio-lateral dissociation in the VOTC (Konkle & Oliva, 2012a). Here, we asked how the neural encoding of real-world size relates to two object properties that are typically covarying with real-world size: shape and spatial mobility. Using human fMRI together with a novel object stimulus set that orthogonalizes real-world size (large, small), shape (rectilinear, curvilinear), and mobility (stationary, mobile), we examined their independent contributions to activity in the PPA, other scene-selective areas, as well as in the visual cortex more broadly.

We report several novel findings: (1) The three object properties each activate the scene-selective PPA, (2) as well as scene-selective OPA and RSC. (3) The three properties cluster specifically in the PPA and OPA, at the intersection of perceptual and memory scene regions. (4) Outside scene-selective regions, the three properties activate divergent regions. Below we discuss each of these findings in more detail.

While the PPA responds strongly to scenes (Epstein & Kanwisher, 1998), studies have shown that it also responds strongly to isolated objects that are large in the real-world (e.g., Aguirre et al., 1998; He et al., 2013; Konkle & Caramazza, 2013; Troiani et al., 2014). However, there is a debate around whether large real-world object responses reflect sensitivity to lower-level visual features that characterize large objects, such as rectilinearity, since these can automatically evoke real-world size knowledge (Long et al., 2017), have an overlapping functional organization as real-world size (Long et al., 2018), and the PPA responds to rectilinear objects and shapes (Nasr et al., 2014; Yue et al., 2020). In the current study, we replicated the PPA’s sensitivity to rectilinear shapes. Interestingly, however, we also found large-object responses in the PPA when controlling rectilinearity, and the size effect was found for both rectilinear and curvilinear objects. Thus, large real-world size responses in the PPA are not reducible to differences in rectilinearity between large and small objects. Instead, the results are in line with the responses reflecting a higher-level process (Luo et al., 2023), possibly related to top-down input to the PPA (Gabay et al., 2016; He et al., 2013; Mullally & Maguire, 2011).

It has been speculated that large objects evoke a higher-level navigationally relevant process in the PPA because of large objects’ suitability for navigation (Aguirre et al., 1998; Epstein & Vass, 2014), consistent with the PPA responding strongly to objects at navigational decision points (vs. non-decision points; Janzen & Van Turennout, 2004; Schinazi & Epstein, 2010). Moreover, damage to the cortical area where the PPA is typically located can lead to impairment in navigation using salient landmarks such as buildings (McCarthy et al., 1996). Large objects may be navigationally relevant because size correlates with being stationary (vs. mobile), a crucial property of landmarks (e.g., Biegler & Morris, 1996). Moreover, studies have shown that stationary (vs. mobile) objects activate the PPA (Mullally & Maguire, 2011; Troiani et al., 2014), although the stationary property was correlated with both large size and rectilinearity in those studies. Notably, we found a preference for large objects in the PPA also when controlling the stationary property, showing that the size effect is not explained by being stationary, and could potentially reflect a conceptual size distinction (Gabay et al. 2016). We also found a stationary effect, when controlling both size and shape, showing that this effect is also not explained by these correlating properties. Interestingly, size and stationary effects interacted, such that the size and stationary effects were smaller for stationary and large objects, respectively. This finding is not in line with the suggestion that the PPA would be particularly responsive to objects that are both large and stationary, owing to these objects being the most space-inducing (Mullally & Maguire, 2011). Indeed, we did not find evidence that the combination of these two properties activated the PPA more than the presumably less space-inducing large-and-mobile objects. Overall, here we find that in addition to low-level rectilinear (vs. curvilinear) properties, objects that are either large or stationary also activate the PPA.

The PPA is part of a larger scene-selective network, including the OPA and RSC (e.g., Epstein & Baker, 2019). Studies have shown that large objects activate OPA and RSC (He et al., 2013), that rectilinear objects activate the OPA (Nasr et al., 2014), and stationary objects RSC (e.g., Auger et al., 2012), but the issue regarding correlation of the properties is equally relevant for the interpretation of these findings as they are for the PPA. Here we find that both cortical regions show a largely similar pattern as the PPA, except for a lack of stationary object preference in RSC. The latter finding is surprising given the RSC’s previous implication in processing landmarks and stationary objects (e.g., Auger et al., 2012). Future studies could test more canonical landmark objects (e.g., buildings) than those included here, while still controlling shape and size, to further examine the role of the RSC in processing landmarks.

A striking finding was that the three properties clustered specifically in and around the PPA and OPA, biased anteriorly to where they overlap with scene memory regions (Steel et al., 2021; 2025). It is striking that the rectilinear effect is present in this more anterior region, also consistent with the finding that the rectilinear effect for real-world objects is more anterior than the effect for simple visual shapes (Nasr et al., 2014). The fact that the properties cluster together in spatial and navigationally related regions could suggest that most objects with these properties are statistically useful for spatial and navigational function, and thus neighboring neurons develop selectivity to these properties based on basic principles of brain organization where neurons with similar functional profiles cluster together spatially (e.g., Aflalo & Graziano, 2011; Durbin & Mitchison, 1990).

The three properties evoked activity in divergent cortical regions outside scene-selective cortex. Large (vs. small) activity was most extensive, located along the medial ventral and dorsal surface, and was strikingly similar to large (vs. small) activity with non-controlled stimuli (**Fig. S2**), and corresponded generally with scene-selective maps (**Fig. S3**). Interestingly, large objects evoked strong activity in ventral posterior regions, consistent with previous findings of conceptual size responses in early visual cortex (Gabay et al., 2016). It could be that the posterior size responses reflect a retinotopic organizational principle, since the regions we find here fall in peripheral view biased visual cortex (Hasson et al., 2003). Additionally, it could reflect feedback from higher-level regions (e.g., Gabay et al., 2016), as observed for size constancy, a mechanism that makes stimuli far away in a scene appear larger than they physically are, an illusion that is reflected in V1 activity (Murray et al., 2006).

We did not find regions responding more strongly to small than large objects for the controlled stimuli, unlike studies showing small-object responses in the lateral VOTC for uncontrolled objects (Konkle & Oliva, 2012a), which we replicated in the uncontrolled contrast (**Fig. S2**). However, in a targeted ROI analysis for small objects, with the ROI located in lateral VOTC (**Fig. S4**), we did find a weak small-object preference in the lateral VOTC for controlled comparisons, together with a curvilinear preference. The curvilinear preference in lateral parts of the VOTC was also observed in the whole-brain contrast, in line with previous work (Yue et al., 2020). The strong preference for small objects in lateral VOTC for the uncontrolled object contrast (Konkle & Oliva, 2012a; Fig. S2) may thus reflect a combination of size and curvilinear preference. Furthermore, previously-observed responses to small objects in lateral VOTC may also have partly reflected action- or body-related processes when perceiving tools and other manipulable objects, objects that are known to drive the lateral VOTC (Cortinovis et al., 2025) and which were not included in the current controlled stimulus set.

Finally, the stationary objects yielded more focal activations, with surprisingly strong stationary activity in lateral occipitotemporal cortex, in the vicinity of the lateral occipital complex (LOC), and mobile activity on the posterior parts of the ventral surface. Given the exploratory nature of this analysis, future work should examine this using a larger object set and more stringent statistical tests.

Overall, the large-scale maps revealed that the properties diverged outside scene-selective cortex, again suggesting that the three property contrasts do not capture one common aspect that was not considered in our design.

In summary, we dissociated object properties that naturally covary in manmade objects and that have all been associated with the functional organization of the human VOTC. After these controls, all properties selectively activated scene-selective regions of visual cortex. These results show that the large-object preference in scene-selective cortex cannot be reduced to rectilinear or stationary object properties. More generally, they show a remarkable convergence of selectivity for three distinct object properties that are highly correlated in naturalistic vision.

## Acknowledgements

European Research Council (ERC) under the European Union’s Horizon 2020 research and innovation programme to M.P. (grant agreement No. 725970). FWO grant 12A4R24N to S.H.

## Data statement

Data will be made available upon publication.

