## Supplemental Info for "Dissociating representations of object shape, real-world size, and mobility in human visual cortex"

**Table S1.**

*Object categories and associated experimental condition.*

| Condition | Object category |
|---|---|
| Large-curvilinear-nonstable | Air balloon |
| Large-curvilinear-nonstable | Car (VW beetle) |
| Large-curvilinear-nonstable | Pool/lake/river float |
| Large-curvilinear-nonstable | Caravan trailer |
| Large-curvilinear-stable | City fountain |
| Large-curvilinear-stable | Garden pool |
| Large-curvilinear-stable | Radar installation |
| Large-curvilinear-stable | Garden trampoline |
| Small-curvilinear-nonstable | Party balloon |
| Small-curvilinear-nonstable | Soccer ball |
| Small-curvilinear-nonstable | Motorcycle helmet |
| Small-curvilinear-nonstable | Couch pillow |
| Small-curvilinear-stable | Bird house |
| Small-curvilinear-stable | Fire alarm |
| Small-curvilinear-stable | Ceiling hanging lamp |
| Small-curvilinear-stable | Waterhouse fixture |
| Large-rectilinear-nonstable | Sailboat |
| Large-rectilinear-nonstable | Cargo trolley |
| Large-rectilinear-nonstable | Car trailer |
| Large-rectilinear-nonstable | Train |
| Large-rectilinear-stable | Bed |
| Large-rectilinear-stable | Building |
| Large-rectilinear-stable | Fridge |
| Large-rectilinear-stable | Shipping container |
| Small-rectilinear-nonstable | Document binder |
| Small-rectilinear-nonstable | Briefcase |
| Small-rectilinear-nonstable | Compost bin |
| Small-rectilinear-nonstable | Kitchen tray |
| Small-rectilinear-stable | Outdoor garden lamp |
| Small-rectilinear-stable | Mailbox |
| Small-rectilinear-stable | Sink |
| Small-rectilinear-stable | Wall mirror |

*Note.* This table shows the composition of object categories in each experimental condition. There were 32 categories across the eight experimental conditions. Each category was represented by three exemplars.

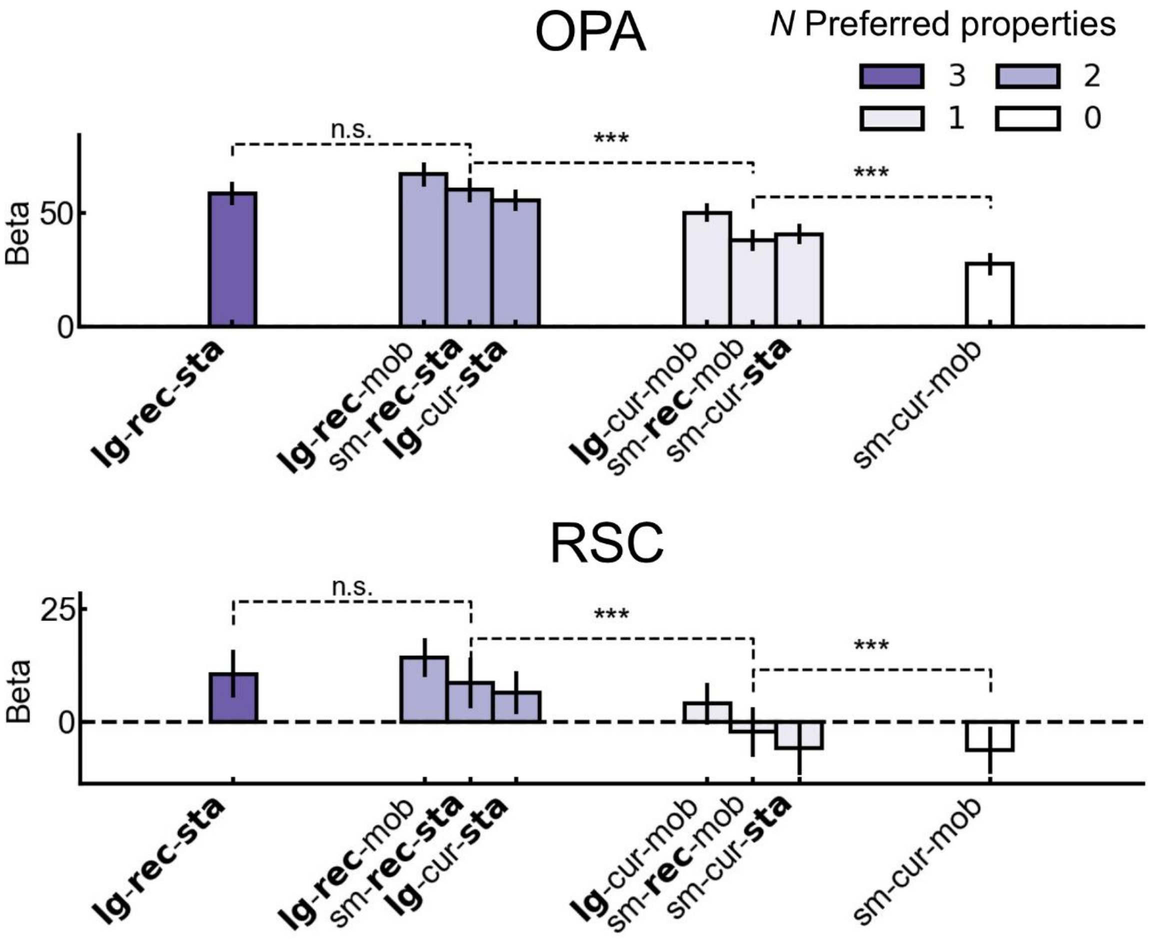


***Figure S1.*** ***Responsiveness of the Occipital Place Area (OPA) and Retrosplenial Complex (RSC) to objects varying on shape (rectilinear, curvilinear), real-world size (large, small), and mobility (stationary, mobile).*** *Beta values for each of the eight experimental conditions, ranked from highest (left) to lowest (right) on preferred properties. Error bars represent 95% CIs. N.s., and *** represent p > 0.05, and p < 0.001, respectively. Error bars represent 95% confidence intervals (CI). Abbreviations: lg: large, sm: small, rec: rectilinear; cur: curvilinear, sta: stationary, mob: mobile.*

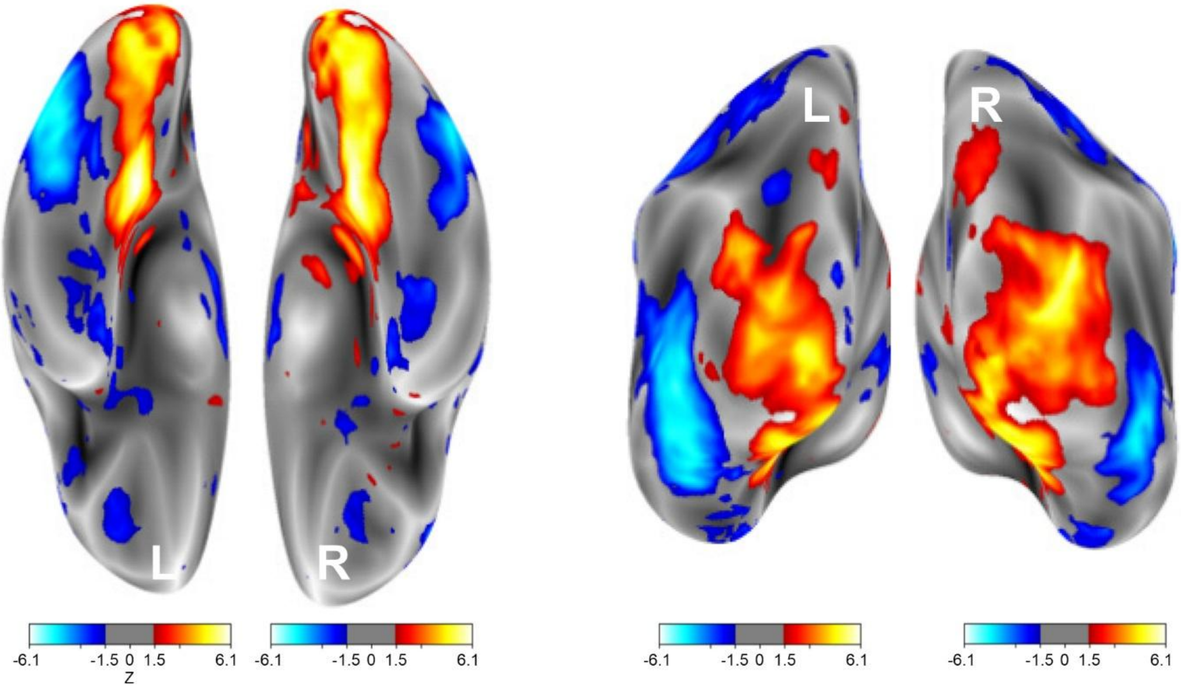


***Figure S2. Group real-world size map from independent localizer data.*** *Localizer stimuli were from Konkle & Oliva (2012), where real-world size covaries with shape and mobility. A liberal threshold of Z = 1.5 is used for visualization purposes.*

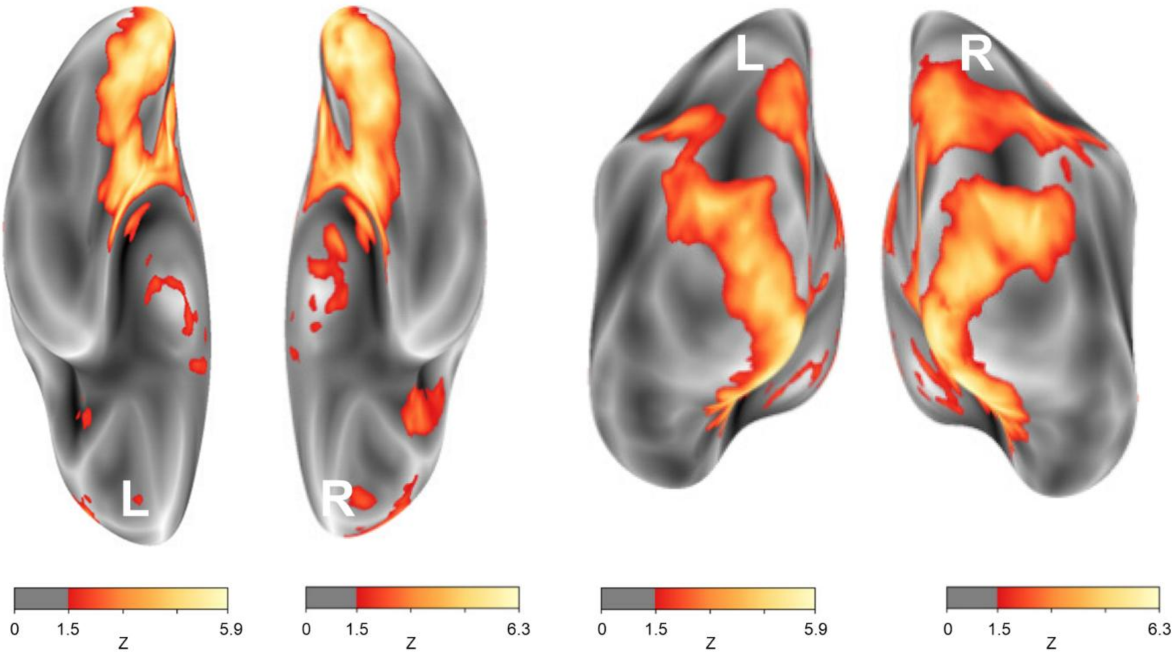


***Figure S3. Group real-world scene-selective map from independent localizer data.*** *Localizer scene-selective maps, where scenes were contrasted with small objects from the*

*Konkle & Oliva (2012) stimulus set. A liberal threshold of Z = 1.5 is used for visualization purposes.*

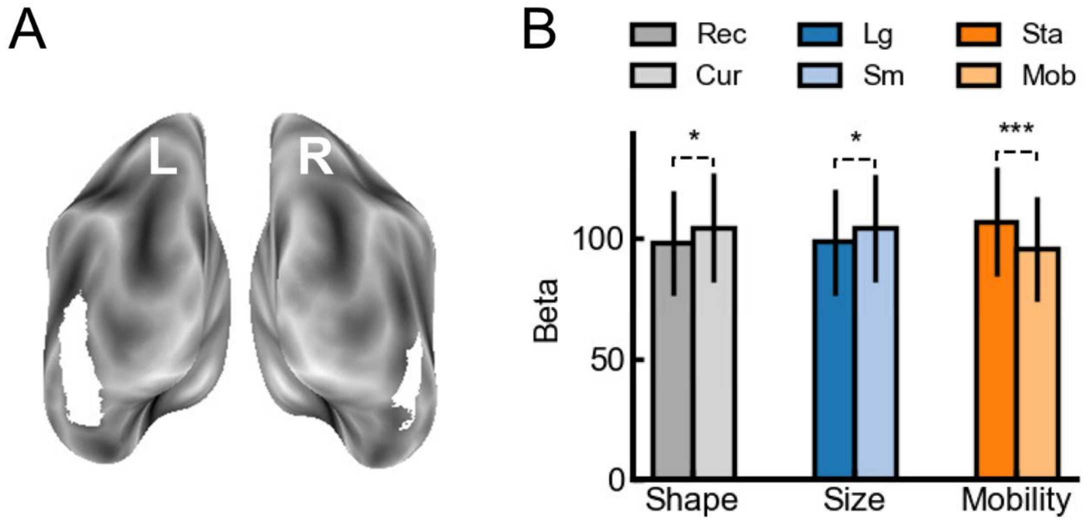


***Figure S4. Responsiveness of the Small (SM) ROI to objects varying in shape (rectilinear, curvilinear), real-world size (large, small), and mobility (stationary, mobile).*** *(**A**) In a region-of-interest (ROI) analysis, a Small ROI was defined by selecting the 50 most small-selective voxels (small > large in independent localizer data) within a bilateral group-level defined Small-ROI mask (from independent localizer data). (**B**) The beta values to shape (rectilinear, curvilinear), size (large, small), and mobility (stationary, mobile) conditions in OPA and RSC. Error bars represent 95% CIs. * and *** represent $p < 0.05$, and $p < 0.001$, respectively.*
